# A Practical Assay for Assessing Mean Red Blood Cell Age Using the Protein 4.1a/4.1b Ratio

**DOI:** 10.64898/2026.10.08.757720

**Authors:** E. Seiler, N. Bender, P. Eppenberger, C. Hobbs-Ray, G. Baerlocher, A. Makhro

## Abstract

**BACKGROUND:** The protein 4.1a/4.1b ratio is an intrinsic molecular marker of mean red blood cell (RBC) age resulting from progressive deamidation of protein 4.1b during RBC circulation. Despite its potential for estimating mean RBC age from a single blood sample, its application has been limited by labor-intensive RBC membrane preparation, and its interindividual variability has not been characterized in a large human population.

**METHODS:** We developed a low-volume method for direct RBC membrane isolation from whole blood using saponin permeabilization and differential centrifugation without prior RBC washing. Membrane proteins were separated by SDS-PAGE, and protein 4.1a/4.1b ratios were determined by densitometric analysis with Gaussian peak fitting. Assay conditions, sample-volume requirements, membrane protein recovery, and analytical reproducibility were evaluated. The method was subsequently applied to 114 anonymized residual samples from routine diagnostic testing selected for complete blood count parameters within the corresponding laboratory reference intervals.

**RESULTS:** The optimized procedure used 50 μL of whole blood, 0.06% saponin, and centrifugation at 9000 g, although protein 4.1a/4.1b remained measurable from starting volumes as low as 5 μL. Mean within-subject CVs were 3.90% for electrophoretic/densitometric determination and 5.39% for membrane protein recovery; whole-procedure CVs in 3 subjects were 1.40%–3.50%. In the population analysis, the mean protein 4.1a/4.1b ratio was 1.428 ± 0.146, with an observed range of 1.052–1.799 and a central 95% interval of 1.151 −1.695. Ratios did not differ significantly between females and males (P=0.246) or among females aged 18 - 50 years, females aged >50 years, and males (P=0.151). The ratio showed modest positive correlations with chronological age in females (Spearman ρ=0.26, P=0.047) and males (ρ=0.33, P=0.015), while substantial interindividual variability remained across the age range.

**CONCLUSIONS:** Direct isolation of RBC membranes from small volumes of whole blood enables reproducible determination of the protein 4.1a/4.1b ratio using a simplified workflow. Application of the assay to a comparatively large human population revealed substantial interindividual variation in this molecular marker of mean RBC age that was not explained by sex or chronological age alone. The method provides a practical approach for investigating biological variation in mean RBC age and its potential contribution to laboratory measurements influenced by RBC survival.

## Introduction

The mean age of circulating red blood cells (RBCs) reflects the balance between erythropoiesis and RBC clearance and provides information that is not captured by conventional measurements of RBC number or hemoglobin concentration **1**. Alterations in RBC production or survival can shift the mean age of the circulating RBC population and affect measurements that accumulate over the RBC lifespan, such as glycated hemoglobin (HbA1c). Indeed, differences in mean RBC age have been shown to produce clinically relevant variation in HbA1c even among hematologically normal individuals **2**. However, established direct methods for determining RBC survival generally require labeling of circulating RBCs or their precursors, followed by serial blood sampling, limiting their applicability in routine laboratory practice **3**,**4**.

Spontaneous protein deamidation provides a naturally occurring molecular clock whose rate is determined by both amino acid sequence and protein structure **5**. Asparagine (Asn) and, more slowly, glutamine (Gln) residues undergo nonenzymatic deamidation, with rates strongly influenced by neighboring amino acids and the local three-dimensional environment; consequently, individual deamidation reactions can have half-times ranging from days to years **5**,**6**. This broad range of intrinsic reaction rates has led to the use of protein deamidation as a molecular timer for biological processes including protein turnover and ageing, and even as an indicator of protein antiquity and preservation in archaeological and palaeontological material **7**,**8**.

Despite being established as an intrinsic marker of RBC age more than three decades ago, the protein 4.1a/4.1b ratio has seen remarkably limited experimental application, having been used in only approximately a dozen research studies, with a substantial contribution from the group of Giampaolo Minetti **9**,**10**,**11**. This limited adoption contrasts with the potential utility of the marker for determining mean RBC age from a single blood sample. One possible practical limitation is the requirement for RBC membrane isolation. Published applications of the protein 4.1a/4.1b ratio have generally relied on conventional RBC ghost preparation, involving RBC purification and washing, hypotonic lysis, and repeated centrifugation and washing steps to remove hemoglobin **9**,**12**. These procedures are based on established methods for preparing hemoglobin-free erythrocyte membranes **13**, while additional purification steps, including cellulose filtration, have been recommended to minimize contamination and preparation-related artefacts **14**. The multiple processing steps required for conventional membrane isolation may therefore represent a practical obstacle to the broader application of the protein 4.1a/4.1b ratio, particularly in routine laboratory settings.

Saponin provides an alternative approach to hypotonic RBC lysis. Through its interaction with membrane cholesterol, saponin increases RBC membrane permeability and induces hemoglobin release while retaining a recoverable membrane structure**15**, **16**. Saponin-permeabilized erythrocytes have consequently been used for decades to investigate membrane transport processes and the activities of membrane-associated enzymes, including Na⁺/K⁺-ATPase and Ca²⁺-ATPase**16**,**17**. We therefore investigated whether saponin permeabilization could be adapted for rapid isolation of RBC membranes suitable for quantitative analysis of protein 4.1a/4.1b.

In the present study, we developed a low-volume assay for direct isolation of RBC membranes from whole blood using saponin permeabilization and differential centrifugation, without prior RBC washing or leukocyte depletion. We optimized the membrane isolation and preanalytical conditions, determined sample-volume requirements, and evaluated the analytical reproducibility of protein 4.1a/4.1b measurement by SDS-PAGE and densitometry. The assay was subsequently applied to residual samples from routine laboratory testing from 114 individuals with complete blood count values within the respective laboratory reference intervals, allowing us to characterize the protein 4.1a/4.1b ratio in a real-world population without laboratory evidence of hematological abnormalities or systemic inflammation. We hypothesized that recurrent menstrual blood loss could influence RBC turnover and reduce mean RBC age in women of reproductive age, whereas women beyond reproductive age would exhibit RBC ages more similar to those of men. To explore this hypothesis, women were divided into two age groups (18–50 and >50 years), while men were analyzed as a single group. Together, these experiments establish a practical approach for assessing mean RBC age from small-volume blood samples and provide a basis for broader research and future clinical application of the protein 4.1a/4.1b marker.

## Methods

### Study samples and ethics

Residual human whole-blood samples were obtained from routine diagnostic laboratory testing at MEDICA Medizinische Laboratorien AG (Zurich, Switzerland) from individuals who had provided general informed consent for the research use of leftover samples from routine diagnostic testing. Samples were provided as fully anonymized biological material and were selected by MEDICA Medizinische Laboratorien AG based on complete blood count parameters and concentrations within the respective laboratory reference intervals. The investigators received fully anonymized samples together with year of birth and sex; individual complete blood count results or other identifying personal data were not provided, and samples could not be traced back to individual subjects. In total, 114 samples meeting the inclusion criteria were included in the population analysis.

The present study was conducted within the framework of the ongoing research project “The role of human Piezo1 mechanosensitive receptor in oxygen delivery efficacy” and was designed on the basis of its established procedure for the use of anonymized human biological material. For this project, the Cantonal Ethics Committee Zurich formally declined jurisdiction (BASEC no. 2024-01985, 4 October 2024), as research involving anonymized biological material without health-related personal data falls outside the scope of the Swiss Human Research Act (HRA, SR 810.30).

### RBC membrane isolation

RBC membranes were isolated directly from whole blood without prior RBC washing or leukocyte depletion. Saponin buffer was prepared freshly before each experiment and consisted of phosphate-buffered saline (PBS) supplemented with 0.06% (w/v) saponin, 1 mmol/L EDTA, and 1 mmol/L phenylmethylsulfonyl fluoride (PMSF). The buffer was maintained on ice throughout the isolation procedure.

Whole blood was thoroughly mixed immediately before sampling. A 50-μL aliquot was added to 1.45 mL of ice-cold saponin buffer in a 1.5-mL microcentrifuge tube and incubated on ice for 2–3 min, until the initially opaque suspension became transparent. Samples were centrifuged for 20 min at 9000 g and 4°C. The red supernatant was aspirated using a vacuum pump equipped with a fine glass tip, leaving approximately 1 mm of liquid above the upper cloudy fraction.

The cloudy fraction was gently rinsed with 200 μL of ice-cold saponin buffer and transferred to a new microcentrifuge tube without disturbing the underlying translucent interface and adherent white-to-pink lower pellet. Ice-cold saponin buffer was added to a final volume of 1.5 mL, and samples were centrifuged for 10 min at 9000 g and 4°C. The supernatant was completely aspirated, leaving the isolated RBC membrane pellet.

For electrophoresis, membrane pellets were mixed with an equal volume of 2× urea/DTT sample buffer to obtain final concentrations of 50 mmol/L Tris-HCl (pH 6.8), 2% (w/v) SDS, 5% (v/v) glycerol, 2 mol/L urea, 1 mmol/L EDTA, 100 mmol/L DTT, and bromophenol blue. Samples were mixed by pipetting, heated at 90°C for 8 min, and briefly centrifuged to collect condensate. Samples not analyzed immediately were stored at −20°C.

### SDS-PAGE and protein staining

RBC membrane proteins were separated by SDS-PAGE using 7.5% polyacrylamide separating gels with a 3% stacking gel in 1-mm-thick Mini-PROTEAN gel cassettes (Bio-Rad Laboratories).

For preparation of 4.6 mL of 7.5% separating gel, 1.15 mL of 4× separating-gel buffer (pH 8.8), 1.15 mL of 30% acrylamide solution, and 2.3 mL of ddH₂O were combined, followed by addition of 19 μL of 8% ammonium persulfate (APS) and 2.2 μL of TEMED. Following polymerization of the separating gel, a 3% stacking gel was added.

Electrophoresis was performed at a constant current of 12 mA during migration through the stacking gel and 15 mA during separation through the 7.5% resolving gel. The total electrophoresis time was approximately 2–2.5 h.

Following electrophoresis, gels were fixed for 15 min in 25% (v/v) isopropanol and 10% (v/v) acetic acid and stained for 30–60 min with 0.006% (w/v) Coomassie Brilliant Blue R-250 in 10% (v/v) ethanol and 10% (v/v) acetic acid. Gels were destained for 40–60 min in 10% (v/v) ethanol and 10% (v/v) acetic acid, with the destaining solution replaced three times.

### Gel imaging and protein 4.1a/4.1b quantification

Destained gels were placed on a uniformly illuminated LED transillumination surface and photographed using a digital camera. Digital images of SDS-PAGE gels were cropped to include the protein 4.1a/4.1b doublet and the protein 4.2 band before densitometric analysis. The cropped images were analyzed using a custom Python script (Supplementary Code 1). The protein 4.2 band was used to identify lane positions and define lane widths, while the protein 4.1a/4.1b doublet was fitted with two Gaussian functions. The protein 4.1a/4.1b ratio was calculated from the background-corrected densitometric profile heights at the fitted Gaussian peak centers.

For each electrophoretic lane, the densitometric profile of the protein 4.1 region was extracted, and the protein 4.1a and 4.1b peaks were identified and fitted using Gaussian functions. The protein 4.1a/4.1b ratio was calculated from the fitted peaks.

For population measurements, each membrane preparation was loaded in five separate electrophoretic lanes, providing five replicate determinations of the protein 4.1a/4.1b ratio for each subject. The mean of the five measurements was used as the subject-level protein 4.1a/4.1b ratio.

### Assay optimization experiments

Assay conditions were evaluated by varying individual experimental parameters while maintaining the remaining conditions unchanged.

Saponin concentrations of 0.02%, 0.04%, 0.06%, and 0.08% were evaluated. Nine independent blood samples were analyzed at each quantifiable concentration. At 0.02% saponin, insufficient protein separation precluded reliable quantification of the protein 4.1a/4.1b ratio; this condition was therefore evaluated qualitatively by SDS-PAGE but was not included in quantitative comparison of the ratios.

The effect of centrifugation speed was evaluated in blood samples from five individuals at 8000, 9000, 10000, 11000, and 14000 g. Aliquots from each individual were processed at each centrifugation speed under otherwise identical conditions and analyzed by SDS-PAGE and protein 4.1a/4.1b densitometry.

Preanalytical stability at room temperature was evaluated using samples from five different individuals. Aliquots were maintained at room temperature (22°C) for up to 196 h before membrane isolation and subsequent SDS-PAGE and protein 4.1a/4.1b analysis.

The effect of cooling before membrane isolation was investigated separately using samples from four individuals. Aliquots were maintained on ice for either 15 or 140 min before processing under otherwise identical conditions.

### Membrane protein recovery and sample-volume experiments

Total protein concentration in isolated RBC membrane preparations was determined by the Bradford assay, using bovine serum albumin (BSA) as the standard and measuring absorbance at 595 nm. Whole-blood hemoglobin concentration was determined by the cyanmethemoglobin method using Drabkin reagent, with absorbance measured at 540 nm and corrected for optical path length.

Total membrane protein recovery from the standard 50-μL whole-blood preparation was determined in samples from 16 different individuals. Membrane protein recovery was expressed both as μg protein per 50 μL of whole blood and, after normalization to the corresponding whole-blood hemoglobin content, as μg protein/mg Hb.

The effect of starting blood volume was investigated using aliquots of 5, 10, 15, 20, 30, and 50 μL obtained from the same blood samples and processed using the same membrane isolation procedure. The resulting preparations were analyzed by SDS-PAGE, and the protein 4.1a/4.1b ratio was determined where sufficient material was available.

### Analytical reproducibility

Reproducibility of the electrophoretic and densitometric determination of the protein 4.1a/4.1b ratio was evaluated using five separately loaded electrophoretic lanes from each study sample. The mean, SD, and coefficient of variation (CV) of the replicate protein 4.1a/4.1b measurements were calculated for each subject.

Consistency and reproducibility of membrane protein recovery were evaluated using blood samples from different individuals. Membrane protein recovery was expressed both as μg protein per 50 μL of whole blood and as μg protein/mg Hb to assess between-subject variability in protein yield before and after normalization to the corresponding whole-blood hemoglobin content. For samples used to assess isolation reproducibility, two independent 50-μL aliquots from the same blood sample were processed separately through the membrane isolation procedure, and membrane protein recovery was determined for each preparation.

Whole-procedure reproducibility was evaluated using blood samples from three individuals. Each blood sample was divided into five independent 50-μL aliquots, which were processed separately through membrane isolation and subsequent SDS-PAGE, gel imaging, and densitometric determination of the protein 4.1a/4.1b ratio, yielding five independently derived protein 4.1a/4.1b results for each subject.

### RBC separation by Percoll density-gradient centrifugation

RBCs were separated into density fractions using a self-forming Percoll gradient. Autologous plasma was first separated from an aliquot of whole blood by centrifugation at 3000 g for 5 min at 30°C and maintained at room temperature for subsequent RBC resuspension. An isotonic Percoll stock solution was prepared by mixing 9 parts Percoll with 1 part 10× phosphate-buffered saline (PBS). A 90% Percoll working solution was then prepared by mixing 9 parts of this stock solution with 1 part 1× PBS.

For density-gradient separation, 1.8 mL of 90% Percoll solution and 200 μL of whole blood were combined in 2-mL microcentrifuge tubes. The tubes were closed, mixed by repeated inversion, and centrifuged at 16 000 g for 35 min at 30°C. Following centrifugation, the upper fraction containing leukocytes was aspirated and discarded. Eight consecutive RBC-containing fractions were then collected from top to bottom by sequential aspiration of 150-μL aliquots. Each fraction was washed three times with PBS to remove residual Percoll (3000 g for 5 min at 30°C per wash) and resuspended in autologous plasma before RBC membrane isolation using the standard saponin-based procedure described above. The density of each RBC-containing fraction was determined gravimetrically using an analytical balance. The procedure was performed in three independent experiments.

### Statistical analysis

For replicate analytical measurements, the arithmetic mean, SD, and coefficient of variation (CV = SD/mean × 100%) were calculated. Technical replicates originating from the same blood sample were not treated as independent biological observations.

Analytical reproducibility of the electrophoretic/densitometric determination of the protein 4.1a/4.1b ratio was assessed from replicate measurements. The pooled within-sample standard deviation was defined as sᵣ = √[Σᵢ(nᵢ − 1)sᵢ² / Σᵢ(nᵢ − 1)], where sᵢ is the standard deviation of replicate measurements for sample i and nᵢ is the number of replicates. The repeatability coefficient was defined as RC = 1.96 × √2 × sᵣ (approximately 2.77sᵣ), representing the expected 95% limit for the absolute difference between two measurements under repeatability conditions. No numerical pooled SD or RC is reported because the replicate-level data are not included in the present manuscript.

To evaluate the number of electrophoretic replicates required for reliable estimation of the protein 4.1a/4.1b ratio, precision was evaluated as a function of the number of replicate measurements included in the reported result. For samples with five replicate measurements, estimates based on one to five replicate measurements were compared with the five-replicate mean, and the reduction in analytical variability with increasing replicate number was assessed.

Membrane protein recovery was evaluated both per fixed volume of whole blood (μg protein/50 μL blood) and after normalization to the corresponding hemoglobin content (μg protein/mg Hb). Between-subject variability in membrane protein recovery was summarized by the mean, SD, and CV for each expression of protein yield. For samples subjected to replicate independent membrane isolations, within-subject isolation reproducibility was assessed from the mean, SD, and CV of protein recovery from the independently processed aliquots.

Whole-procedure reproducibility of the protein 4.1a/4.1b measurement was assessed separately for each of the three subjects using five independently processed aliquots. The mean, SD, and CV of the five independently derived protein 4.1a/4.1b ratios were calculated for each subject.

Assay-optimization experiments were analyzed as repeated-measures experiments when the same blood samples were evaluated under multiple experimental conditions. Protein 4.1a/4.1b ratios obtained with 0.04%, 0.06%, and 0.08% saponin were compared using the Friedman test. The 0.02% saponin condition was excluded from the quantitative comparison because membrane protein separation was insufficient for reliable determination of the protein 4.1a/4.1b ratio in all samples. Centrifugation speeds were similarly compared using the Friedman test because the same samples were evaluated at each speed. Preanalytical room-temperature storage was evaluated from paired measurements obtained from the same samples before and after storage and was summarized descriptively because of the small number of samples.

For population analysis, the mean of the five replicate protein 4.1a/4.1b measurements was used as a single subject-level value. The distribution of subject-level protein 4.1a/4.1b ratios was summarized using descriptive statistics and examined graphically. A central 95% distribution interval was estimated from the 2.5th and 97.5th percentiles of the subject-level distribution. Associations between the protein 4.1a/4.1b ratio and age were assessed using Spearman rank correlation. Differences in protein 4.1a/4.1b ratios between demographic groups were evaluated using statistical tests appropriate to the final group structure and distribution of the data. All statistical tests were two-sided, and *P* < 0.05 was considered statistically significant.

## Results

### 1. Development of a rapid whole-blood membrane isolation assay

To facilitate routine determination of the protein 4.1a/4.1b ratio, we developed a rapid membrane isolation assay requiring only 50 μL of whole blood, a volume compatible with finger-prick blood collection. Following saponin permeabilization and centrifugation, the sample reproducibly separated into an upper cloudy membrane fraction, a thin translucent interface, and a lower white pellet (Fig. 1A). Selective recovery of the upper fraction, rather than the entire pellet, provided clean membrane preparations suitable for SDS-PAGE while avoiding material from the interface and lower pellet.

**Figure 1.**
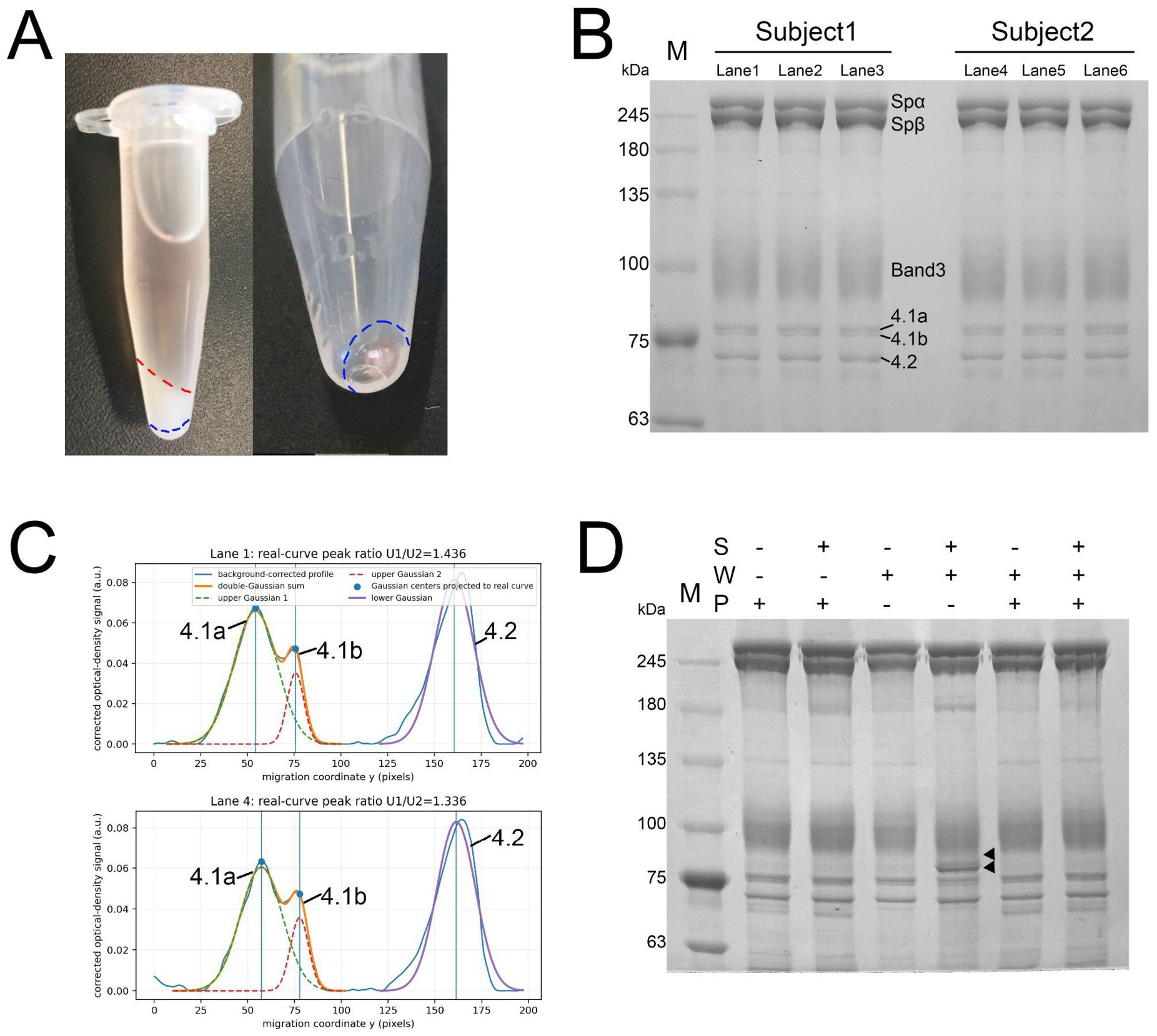
Whole-blood RBC membrane isolation assay and representative readout. **(A)** Following saponin permeabilization and centrifugation, the sample separates into an upper cloudy membrane fraction, a thin translucent interface, and a lower white pellet. The upper fraction between the red and blue dashed lines was selectively collected for analysis, leaving the interface and lower pellet undisturbed. The right image shows the isolated membrane pellet after the subsequent washing and centrifugation step. **(B)** Representative separation of isolated RBC membranes on a 7.5% SDS-polyacrylamide gel. Major RBC membrane proteins, including α- and β-spectrin, band 3, proteins 4.1a/4.1b, and protein 4.2, are resolved. Protein 4.1b and its deamidated form, protein 4.1a, are indicated. Three replicate lanes from each of two individuals illustrate differences in the relative abundance of protein 4.1a and 4.1b. **(C)** Representative densitometric analysis of lanes 1 and 4 from panel B (see Methods). The heights of the fitted protein 4.1a and 4.1b peaks were used to calculate the protein 4.1a/4.1b ratio. The ratios were 1.436 for lane 1 (Subject 1) and 1.336 for lane 4 (Subject 2). **(D)** Effect of RBC washing and plasma reconstitution on the membrane protein pattern. Whole blood and washed RBCs were incubated for 60 min at 37°C in the absence of EDTA and PMSF. S indicates the presence (+) or absence (−) of 0.06% saponin, W indicates three consecutive RBC washing steps, and P indicates the presence of autologous plasma. Additional lower-molecular-weight bands within the resolved range (≥63 kDa), indicated by black arrowheads, appeared in washed RBC preparations in the absence of plasma and likely represent proteolytic products of high-molecular-weight membrane proteins. Resuspension of washed RBCs in autologous plasma prevented the appearance of these additional bands.

Electrophoretic separation consistently resolved the major RBC membrane proteins, including the closely migrating protein 4.1a and 4.1b bands required for determination of the protein 4.1a/4.1b ratio (Fig. 1B). Representative membrane preparations from two individuals illustrate differences in the relative abundance of protein 4.1a and 4.1b and consequently in the protein 4.1a/4.1b ratio. The ratio was determined by densitometric analysis of the corresponding electrophoretic bands (Fig. 1C). Protein 4.1a is generated by age-dependent deamidation of protein 4.1b, making the protein 4.1a/4.1b ratio a surrogate marker of the mean age of circulating RBCs.

The effect of prior RBC washing was examined under conditions favoring proteolysis. Following incubation with saponin for 60 min at 37°C in the absence of EDTA and PMSF, washed RBCs showed additional lower-molecular-weight protein bands within the range resolved on the gels (≥63 kDa), whereas these changes were not observed in membrane preparations obtained directly from whole blood (Fig. 1D). In a reconstruction experiment, washed RBCs were resuspended in autologous plasma before incubation under the same conditions; the additional bands were no longer observed, and the electrophoretic pattern was comparable to that obtained from whole blood. Based on these observations, RBC washing was omitted from the final membrane isolation procedure.

### 2. Optimization of membrane isolation and pre-analytical conditions

To establish robust assay conditions for routine laboratory use, we optimized the saponin concentration, centrifugation speed, and sample handling prior to membrane isolation (Fig. 2).

**Figure 2.**
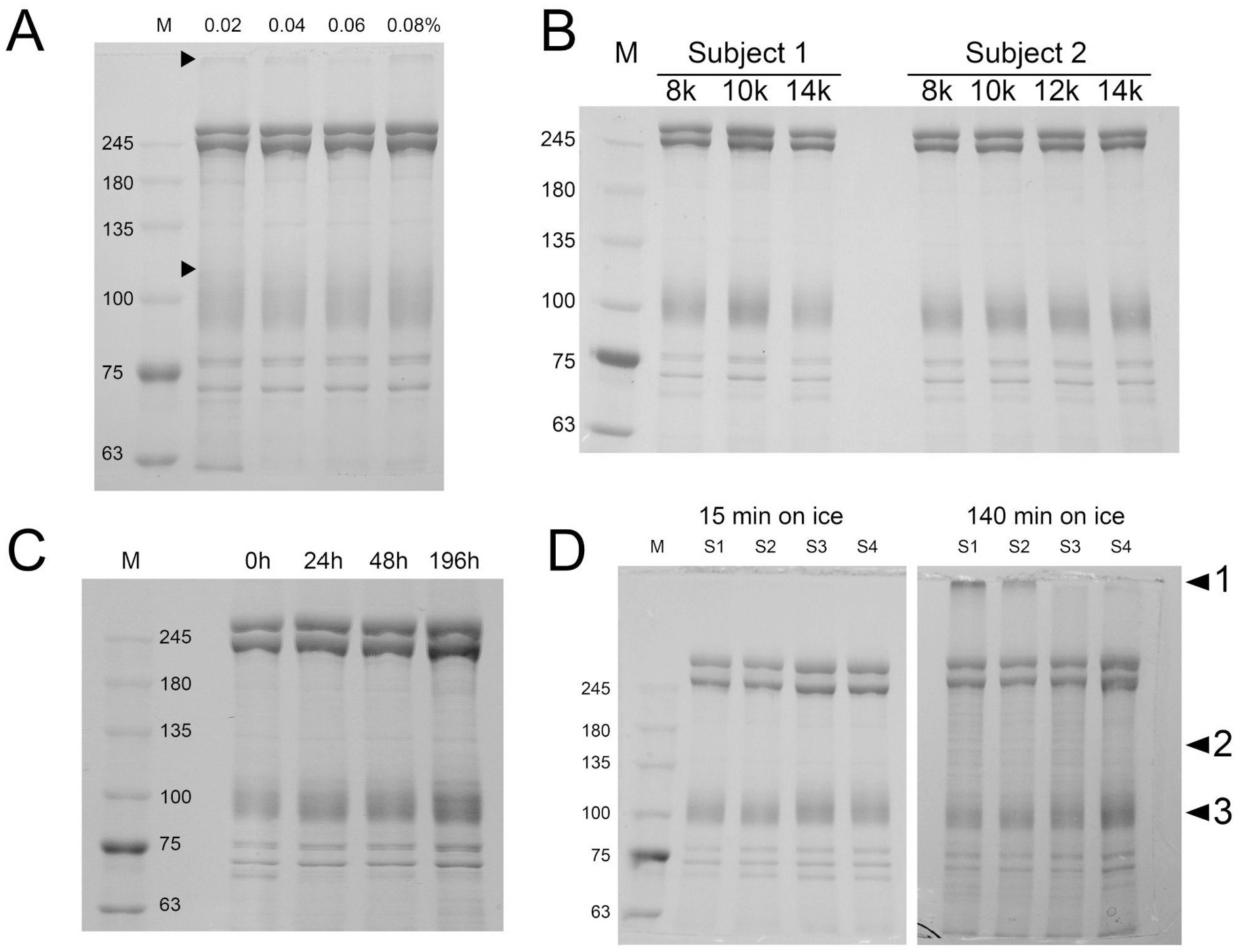
Optimization of membrane isolation and pre-analytical conditions. **(A)** Effect of saponin concentration on RBC membrane isolation and electrophoretic protein separation. Membranes were isolated from the same blood donor using 0.02–0.08% saponin. The lower arrow indicates blurred protein separation observed at 0.02% saponin, whereas the upper arrow marks protein accumulation at the stacking–separating gel interface. **(B)** Effect of centrifugation speed on membrane isolation. Representative membrane preparations obtained at centrifugation speeds between 8,000 g and 14,000 g from two blood donors are shown. No apparent differences in membrane protein patterns were observed. Analysis of five independent donors revealed no statistically significant differences in the protein 4.1a/4.1b ratio between centrifugation conditions. **(C)** Effect of room-temperature storage before membrane isolation in two subjects. Representative membrane preparations obtained after storage of whole blood for up to 196 h at room temperature. No apparent deterioration of the membrane protein pattern was observed. **(D)** Effect of prolonged sample cooling before membrane isolation. Storage of whole blood on ice for 140 min impaired membrane isolation. Arrow 1 indicates protein accumulation at the stacking–separating gel interface; arrow 2, diffuse protein staining; arrow 3, reduced resolution of band 3.

Saponin concentrations from 0.02% to 0.08% were evaluated for membrane isolation from 50 μL of whole blood (Fig. 2A). At 0.02%, membrane preparations showed blurred electrophoretic protein separation, poor resolution of the protein 4.1a/4.1b doublet, and retention of protein material at the stacking–separating gel interface. Although 0.04% produced acceptable membrane preparations in most samples, reduced protein separation was occasionally observed, particularly in older donors. In contrast, 0.06% consistently yielded high-quality protein separation with clear resolution of the protein 4.1a and 4.1b bands. At 0.08%, protein retention at the stacking–separating gel interface was again observed. Therefore, 0.06% saponin was selected for all subsequent experiments.

Centrifugation speeds between 8000 g and 14000 g all produced the characteristic three-component pellet and preserved the RBC membrane protein pattern (Fig. 2B). No statistically significant differences in the protein 4.1a/4.1b ratio were detected in five independent donors. However, lower centrifugation speeds produced a larger visible RBC membrane fraction, facilitating membrane collection while maintaining separation from the lower fractions. Therefore, 9000 g was selected for the final protocol.

Whole blood stored at room temperature for up to 196 h showed no visible deterioration of the membrane protein pattern (Fig. 2C). The protein 4.1a/4.1b ratio remained stable over this period, with mean values of 1.524 immediately after blood collection and 1.513 after 196 h of storage (Table 1).

**Table 1.**
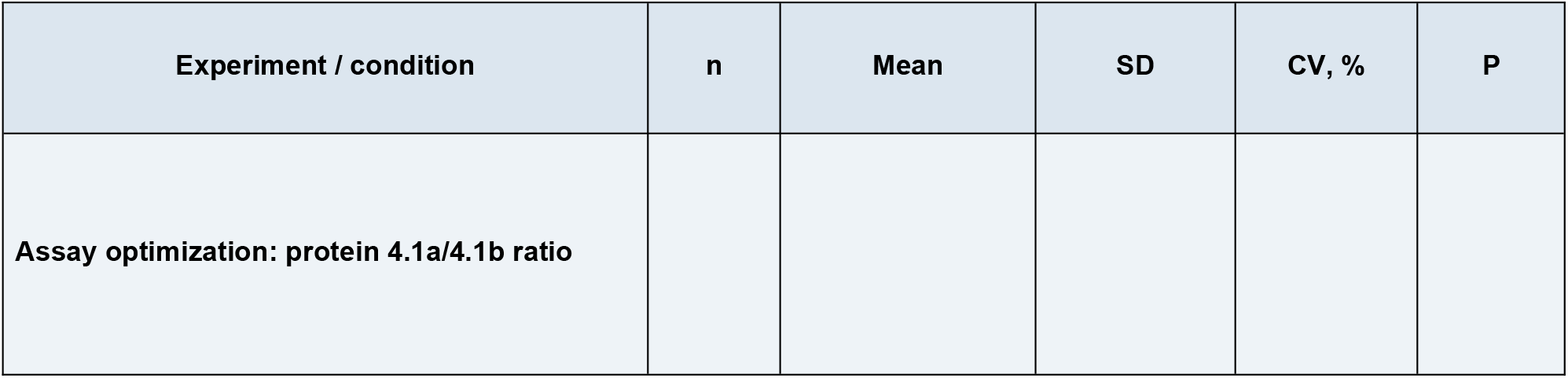
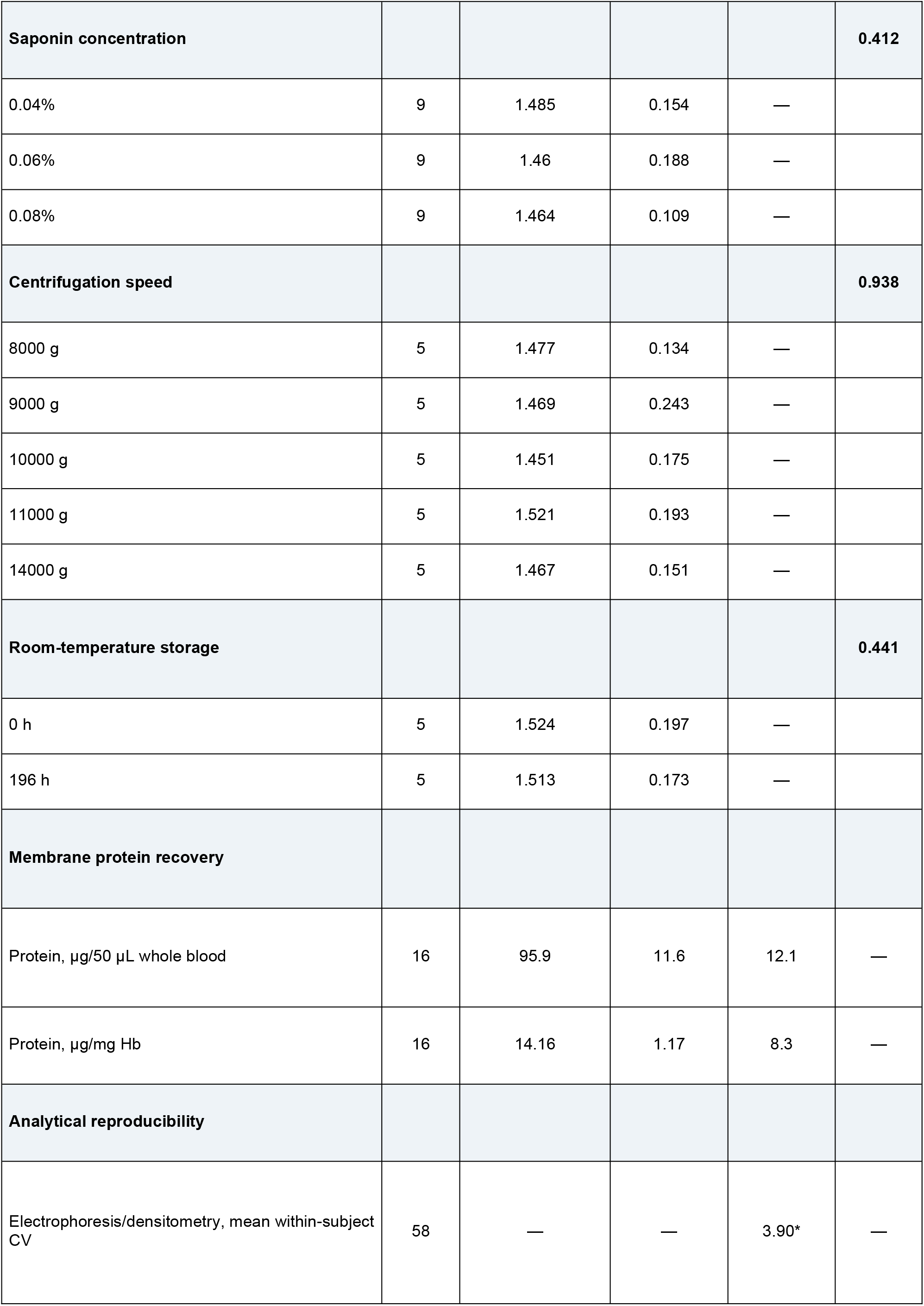
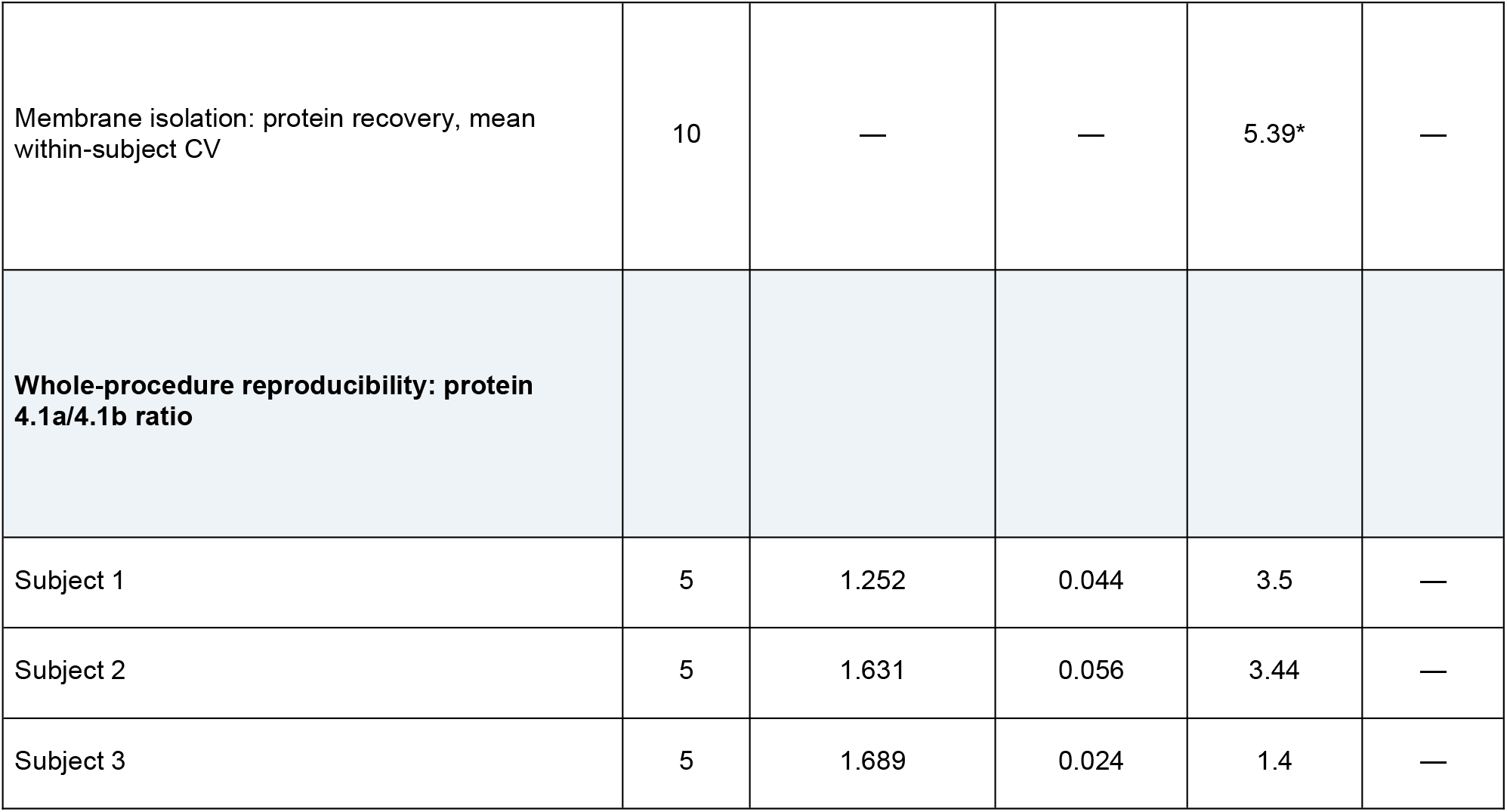
Assay optimization, membrane protein recovery, and analytical reproducibility. Values for assay-optimization experiments represent the protein 4.1a/4.1b ratio. Saponin concentrations of 0.04%, 0.06%, and 0.08% were evaluated in the same 9 subjects and compared using the Friedman test; the 0.02% condition was excluded from quantitative comparison because reliable determination of the protein 4.1a/4.1b ratio was not possible in all samples. Centrifugation speeds from 8000 to 14000 g were evaluated in 5 subjects and compared using the Friedman test. Room-temperature storage was evaluated in 5 subjects by paired measurements at 0 and 196 h. Membrane protein recovery from 50 μL of whole blood was evaluated in 16 subjects and expressed both per fixed blood volume and after normalization to whole-blood Hb. Reproducibility of membrane isolation was evaluated in 10 subjects using two independently processed 50-μL aliquots from each blood sample. Electrophoretic/densitometric reproducibility was evaluated from replicate protein 4.1a/4.1b measurements within individual samples. Whole-procedure reproducibility was evaluated in three subjects using five independently processed 50-μL aliquots per subject. *Mean of subject-specific within-subject CVs. CV, coefficient of variation; Hb, hemoglobin.

In contrast, storage of whole blood on ice for more than 2 h markedly impaired membrane isolation (Fig. 2D), resulting in protein retention at the stacking–separating gel interface, diffuse electrophoretic protein patterns, and reduced resolution of band 3. Therefore, whole blood should be stored at room temperature before membrane isolation.

Based on these experiments, the final protocol consisted of membrane isolation from 50 μL of whole blood using 1.45 mL of 0.06% saponin, centrifugation at 9000 g, and room-temperature storage before membrane isolation.

### 3. Analytical performance and sample requirements

Membrane protein recovery from 50 μL of whole blood was evaluated in 16 subjects. The mean protein yield was 95.9 ± 11.6 μg per 50 μL of whole blood. After normalization to the corresponding whole-blood Hb concentration, the mean recovery was 14.16 ± 1.17 μg protein/mg Hb (Table 1). Between-subject variability was lower when membrane protein recovery was normalized to Hb (CV, 8.3%) than when expressed per fixed blood volume (CV, 12.1%).

Reproducibility of membrane isolation was evaluated separately in 10 subjects using two independently processed 50-μL aliquots from each blood sample. The mean within-subject CV of membrane protein recovery was 5.39% (Table 1).

To determine the minimum blood volume required for analysis, membrane preparations were generated from 5, 10, 15, 20, 30, and 50 μL aliquots from four subjects. Major membrane proteins, including the protein 4.1a and 4.1b bands, were resolved across the complete range of starting volumes, including preparations obtained from 5 μL of whole blood (Fig. 3). No significant differences in the protein 4.1a/4.1b ratio were observed across the tested sample volumes (data not shown).

**Figure 3.**
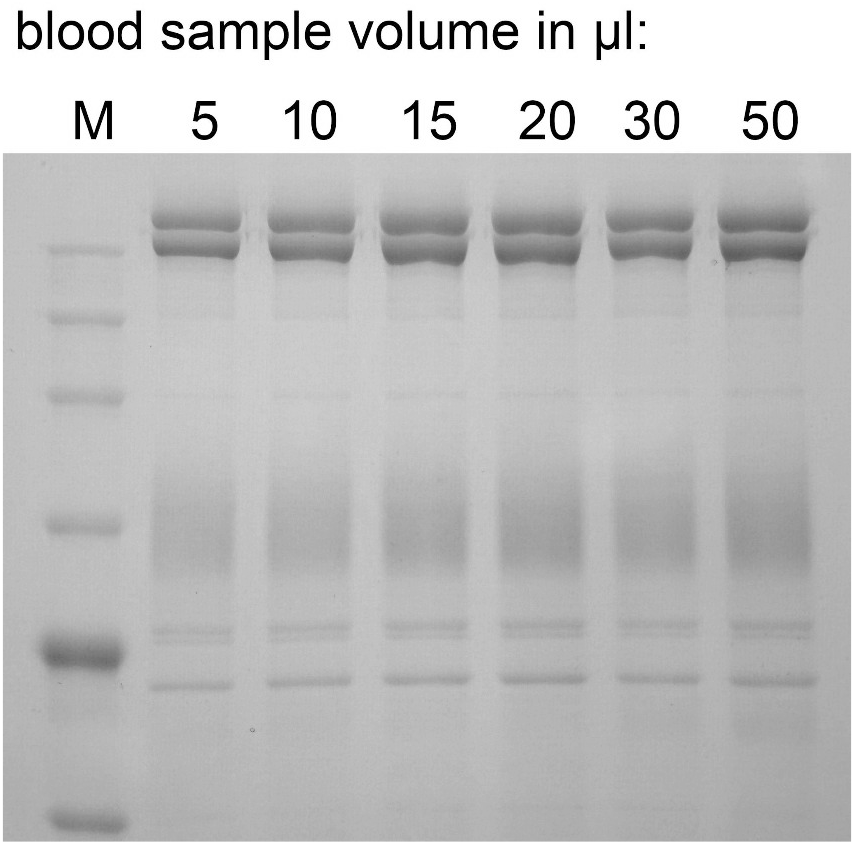
Effect of starting blood volume on RBC membrane protein analysis. Representative SDS-PAGE analysis of membrane preparations isolated from 5, 10, 15, 20, 30, and 50 μL aliquots of the same whole-blood sample. Major RBC membrane proteins, including the protein 4.1a/4.1b doublet, were resolved across the complete range of starting volumes. The experiment was performed using blood samples from four subjects. No significant differences in the protein 4.1a/4.1b ratio were observed across the tested starting volumes (data not shown). M, molecular-mass marker.

Whole-procedure reproducibility was evaluated in three subjects using five independently processed 50-μL aliquots from each blood sample. The mean protein 4.1a/4.1b ratios were 1.252, 1.631, and 1.689, with corresponding within-subject CVs of 3.50%, 3.44%, and 1.40%, respectively (Table 1).

### 4. Detection of protein 4.1a/4.1b differences in density-separated RBC fractions

To evaluate whether the assay could detect differences in the protein 4.1a/4.1b ratio between RBC subpopulations, erythrocytes were separated into eight fractions using self-forming Percoll density-gradient centrifugation. The mean densities of fractions 1–8, collected from top to bottom, were 1.0810, 1.0876, 1.0880, 1.0902, 1.0928, 1.0964, 1.1072, and 1.1104 g/mL, respectively. The approximate positions of the collected fractions are illustrated in Fig. 4A.

**Figure 4.**
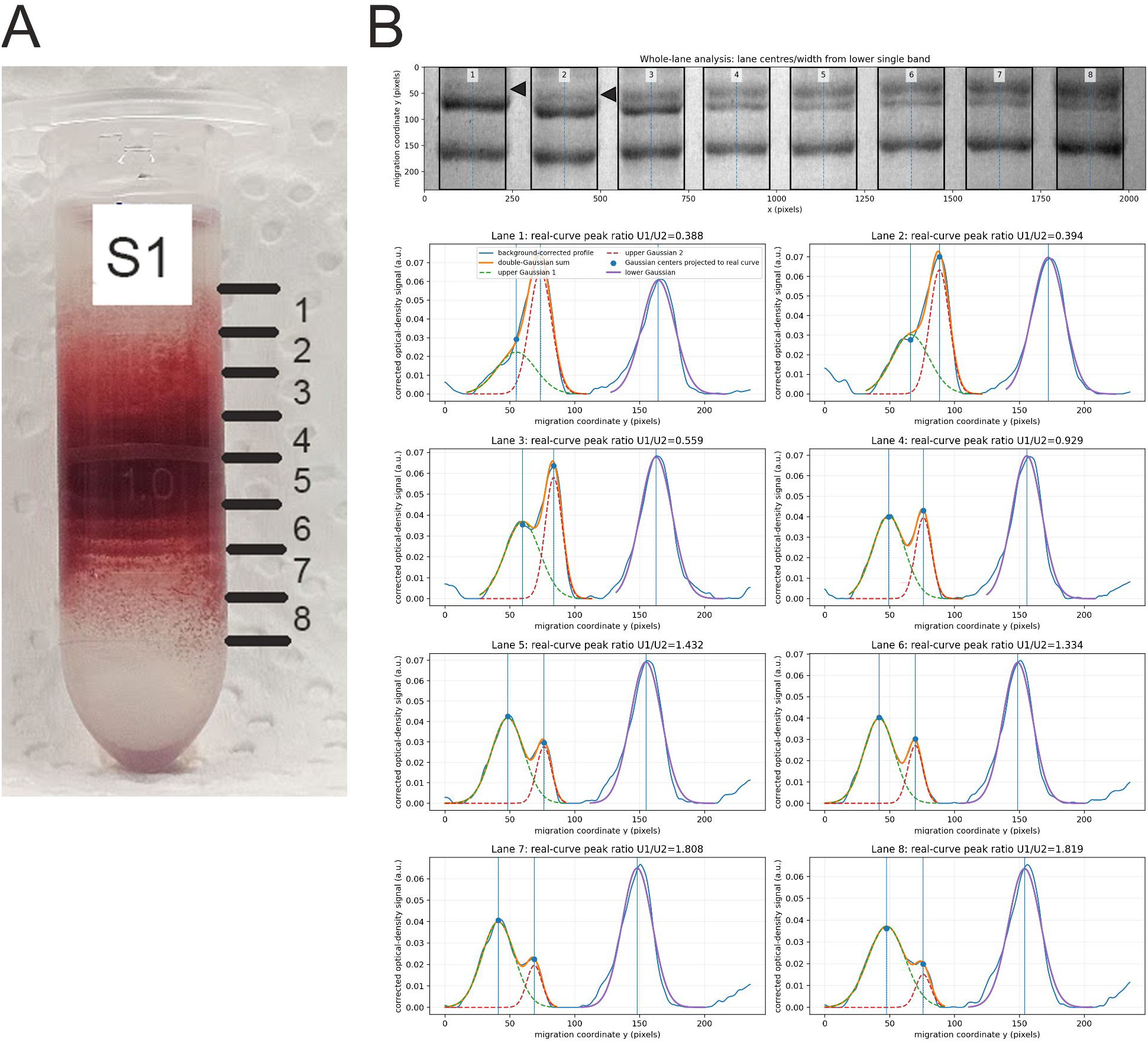
Protein 4.1a/4.1b ratios in RBC fractions separated by Percoll density-gradient centrifugation. (A) Representative separation of erythrocytes into eight density fractions using a self-forming Percoll gradient. Fractions were collected sequentially from top (1, lowest density) to bottom (8, highest density). (B) SDS-PAGE analysis of RBC membrane proteins isolated from the eight fractions (upper panel) and corresponding densitometric profiles of the protein 4.1a/4.1b doublet (lower panels). Protein 4.1a (green) and protein 4.1b (red) peaks were fitted using Gaussian functions, and their peak-height ratios were used to calculate the protein 4.1a/4.1b ratio. The ratio increased from 0.388 in the lowest-density fraction to 1.819 in the highest-density fraction, consistent with progressive enrichment of older RBCs in denser fractions. Representative results from one of three independent experiments are shown.

Representative results from one of three independent experiments are shown in Fig. 4B. The lowest-density fractions exhibited a prominent protein 4.1b band and a comparatively weak protein 4.1a band, as demonstrated by Gaussian fitting of the corresponding densitometric peaks. With increasing RBC fraction density, the relative abundance of protein 4.1a progressively increased, becoming greater than that of protein 4.1b in the densest fraction, where the protein 4.1a/4.1b ratio reached 1.819. These findings demonstrate that the assay can detect differences in the protein 4.1a/4.1b ratio across RBC density fractions, consistent with the expected enrichment of older erythrocytes in higher-density fractions.

### 5. Application of the assay to routine clinical samples

The assay was applied to 114 leftover blood samples originally collected for routine diagnostic testing and selected based on complete blood count parameters and C-reactive protein concentrations within the respective laboratory reference intervals. For samples with replicate protein 4.1a/4.1b measurements, the mean of the available measurements was used as the subject-level value. Across the study population, the mean protein 4.1a/4.1b ratio was 1.428 ± 0.146, with a median of 1.430 (IQR, 1.343–1.531) and an observed range of 1.052–1.799. The central 95% of observed values ranged from 1.151 to 1.695.

The protein 4.1a/4.1b ratio did not differ significantly between females (n=61) and males (n=53; Mann –Whitney U test, P=0.246; Fig. 5A). Among females, the median ratio was 1.386 (IQR, 1.317–1.457) in those aged 18–50 years (n=34) and 1.446 (IQR, 1.351–1.586) in those aged >50 years (n=27). Comparison of females aged 18–50 years, females aged >50 years, and males showed no significant overall difference (Kruskal–Wallis test, P=0.151; Fig. 5B). Substantial interindividual variation in the protein 4.1a/4.1b ratio was observed within both sexes and within each female age group.

**Figure 5.**
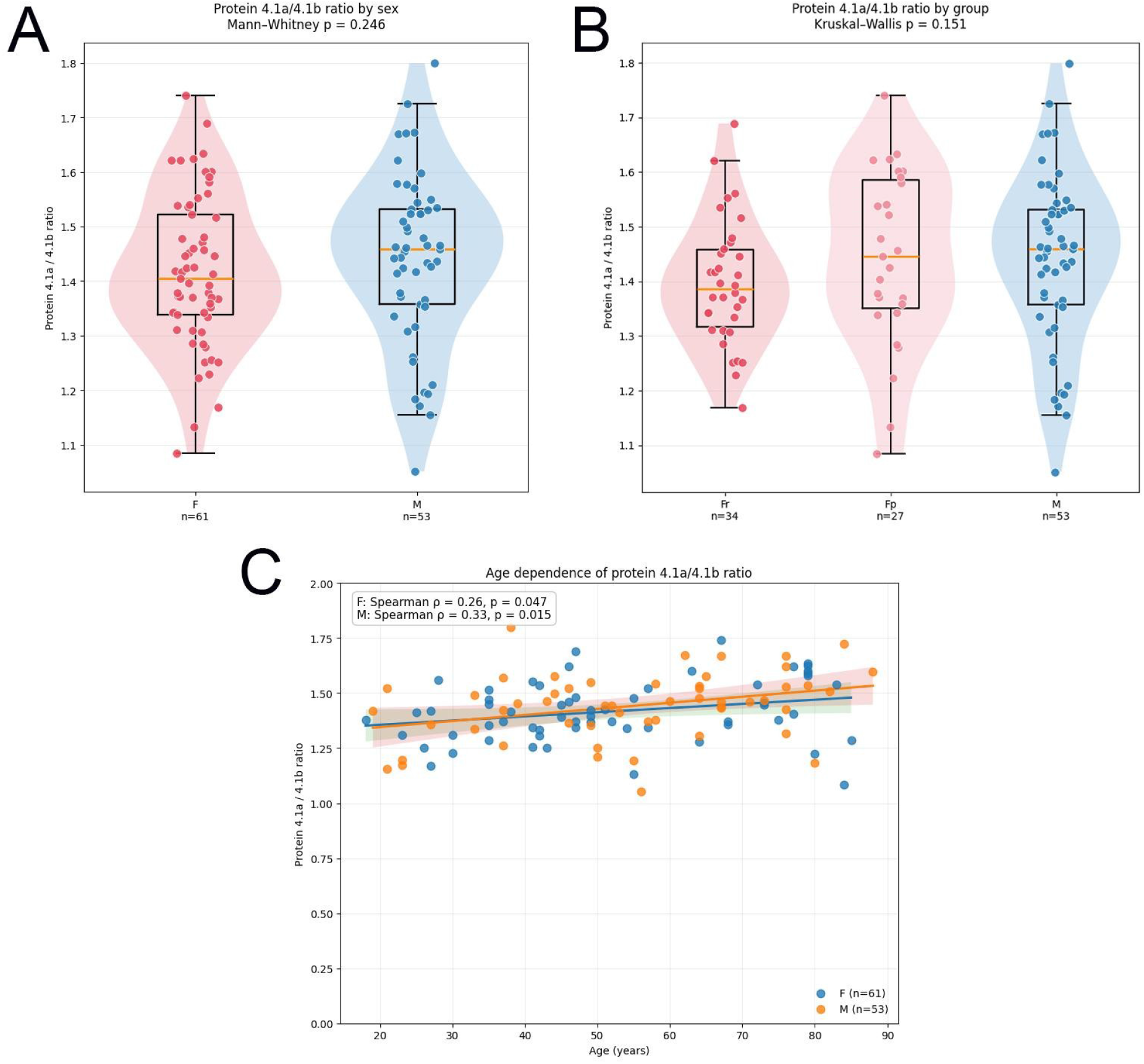
Distribution of the protein 4.1a/4.1b ratio according to sex and age in routine clinical samples. **(A)** Distribution of subject-level protein 4.1a/4.1b ratios in females (F, n=61) and males (M, n=53). Individual values are shown together with violin plots and box plots; orange lines indicate group medians. Groups were compared using the Mann–Whitney U test (P=0.246). **(B)** Protein 4.1a/4.1b ratios in females aged 18–50 years (Fr, n=34), females aged >50 years (Fp, n=27), and males (M, n=53). Groups were compared using the Kruskal–Wallis test (P=0.151). **(C)** Association between protein 4.1a/4.1b ratio and chronological age in females and males. Spearman rank correlations were ρ=0.26 (P=0.047) for females and ρ=0.33 (P=0.015) for males. Lines show linear fits with 95% confidence intervals for visualization. Substantial interindividual variation in the protein 4.1a/4.1b ratio was observed within both sexes and across the investigated age range.

The protein 4.1a/4.1b ratio showed a modest positive association with chronological age in both sexes (Fig. 5C). In females, the ratio correlated with age (Spearman ρ=0.26, P=0.047); in males, the corresponding correlation was ρ=0.33 (P=0.015). Despite these age associations, substantial interindividual variation remained across the entire age range in both females and males.

## Discussion

In this study, we developed a low-volume method for direct isolation of RBC membranes from whole blood and determination of the protein 4.1a/4.1b ratio, an established molecular marker associated with mean RBC age. The method omits prior RBC washing and requires only small volumes of whole blood, while providing reproducible separation of the protein 4.1a/4.1b doublet. The assay was robust across the tested saponin concentrations and centrifugation conditions, and membrane proteins could be analyzed from starting volumes as low as 5 μL. Reproducibility experiments showed low analytical variability, whereas application of the assay to routine diagnostic samples revealed substantial interindividual variation in the protein 4.1a/4.1b ratio.

The protein 4.1a/4.1b ratio has been recognized as an intrinsic marker of RBC age for more than three decades**18**,**19**. Protein 4.1b is progressively converted to protein 4.1a during RBC aging, and this conversion has subsequently been attributed to deamidation, providing a molecular clock of RBC age**9**,**20**. Despite this, the protein 4.1a/4.1b ratio has been used in relatively few studies. A practical limitation may be the labor-intensive preparation of RBC membranes. Conventional methods involve separation and repeated washing of erythrocytes, often including cellulose filtration to remove leukocytes and platelets, as recommended by Minetti and colleagues **14**, followed by hypotonic lysis and repeated membrane washing according to protocols derived from the classical method of Dodge et al. **13**. Our results demonstrate that these preliminary purification steps are not required for reliable resolution and densitometric quantification of protein 4.1a and 4.1b. Moreover, under conditions favoring proteolysis, washed RBC preparations developed additional lower-molecular-weight bands, whereas these changes were prevented by reconstitution with autologous plasma. Although the underlying mechanism was not investigated, these findings indicate that direct processing of whole blood not only simplifies membrane preparation but may also reduce alterations introduced during sample handling.

The method showed encouraging analytical robustness. Protein 4.1a/4.1b ratios were not significantly affected by the tested saponin concentrations or centrifugation speeds, although 0.06% saponin and 9000 g provided the most consistently suitable membrane preparations. Prolonged storage at room temperature produced little change in the ratio in the samples examined, whereas prolonged cooling before membrane isolation adversely affected electrophoretic separation. In addition, the protein 4.1a/4.1b doublet remained measurable across starting blood volumes from 5 to 50 μL. This low sample-volume requirement may be advantageous when sample availability is limited and could potentially allow adaptation of the assay to capillary blood collection, although capillary samples were not investigated in the present study.

Previous studies employing the protein 4.1a/4.1b ratio have generally been based on small numbers of individuals or have used the ratio as an RBC-age marker in experimental settings rather than investigating its natural interindividual variability. Inaba and Maede reported protein 4.1a/4.1b ratios in only four human subjects **19**, while Ravindranath et al. examined seven children with transient erythroblastopenia **21**. Subsequent studies, including work by Minetti and colleagues, primarily used the protein 4.1a/4.1b ratio to characterize RBCs separated into age- or density-related fractions, often from only a few blood donors **9**. In our previous study, the ratio was measured longitudinally in 12 healthy male subjects during and after high-altitude exposure **22**. Thus, although the protein 4.1a/4.1b ratio has been used experimentally as an RBC-age marker for several decades, to our knowledge the present study, comprising 114 individuals, is the first to examine its distribution in a comparatively large human population and to specifically assess its natural interindividual variability.

The ability of the simplified assay to detect age-associated differences in RBC membrane protein composition was further examined using Percoll density-gradient fractionation. Previous studies have demonstrated that the protein 4.1a/4.1b ratio generally increases with RBC density, reflecting the preferential enrichment of older erythrocytes in denser fractions **23**,**12**. Consistent with these observations, we found an overall increase in the protein 4.1a/4.1b ratio across eight density-separated RBC fractions, from 0.388 in the least dense fraction to 1.819 in the densest fraction. This pronounced shift, from predominantly protein 4.1b to predominantly protein 4.1a, demonstrates that the simplified membrane isolation and densitometric procedure can resolve the expected differences between RBC subpopulations. Although RBC density is not an absolute indicator of cellular age, these findings provide additional support for the biological relevance of the measurements obtained with the assay.

Importantly, the magnitude of between-subject variation in the protein 4.1a/4.1b ratio substantially exceeded the analytical variability observed in the reproducibility experiments. No significant difference was observed between females and males or among females aged 18–50 years, females aged >50 years, and males. Nevertheless, considerable variation was present within each group. The ratio showed a modest positive association with chronological age in both females and males, but individuals of similar chronological age frequently had markedly different protein 4.1a/4.1b ratios. Thus, sex and chronological age explained only a limited proportion of the observed population variability.

Applying the tentative regression relationship between the protein 4.1a/4.1b ratio and erythrocyte life span proposed by Inaba and Maede **19**, the ratios observed in the present study corresponded to an estimated mean erythrocyte life span of 121.3 days, with individual estimates ranging from 85.7 to 169.3 days. Mean estimates were 119.9 days in females and 122.9 days in males, and 117.4 and 123.2 days in females aged 18–50 and >50 years, respectively. These values are consistent with the approximately 120-day life span conventionally attributed to human erythrocytes. However, the Inaba equation was derived from an interspecies regression rather than calibrated against individual human RBC survival measurements and should therefore be regarded as an illustrative conversion rather than a validated measurement of RBC life span **19**. In the original study, the authors themselves described the equation as a tentative regression formula.

This finding is particularly relevant to the interpretation of protein 4.1a/4.1b as a marker of RBC age. Chronological age of the individual and mean age of the circulating RBC population are distinct biological variables. Mean RBC age depends on the balance between RBC production and removal and may therefore differ between individuals of the same chronological age. Indeed, substantial variation in RBC survival and mean RBC age has been demonstrated even among apparently healthy individuals **24**. The relatively small analytical variability of the present assay compared with the broad population distribution suggests that a substantial component of the observed variation is biological rather than analytical. Whether this represents stable differences between individuals or temporal variation within individuals remains unknown. Repeated measurements in the same subjects will therefore be important for establishing within-person biological variation of the protein 4.1a/4.1b ratio.

Such interindividual differences in RBC age may be relevant to laboratory measurements that depend on the duration of RBC circulation. HbA1c provides a prominent example, because its concentration reflects both glycemic exposure and the time available for hemoglobin glycation. Variation in RBC survival can consequently contribute to differences between HbA1c and other measures of glycemia even in individuals without overt hematological abnormalities **2**,**24**,**25**. A single-sample marker of mean RBC age could potentially help distinguish variation related to RBC population age from variation related to glycemic exposure. This possibility remains to be tested by directly comparing the protein 4.1a/4.1b ratio with independent measures of RBC survival and with laboratory parameters known to be influenced by RBC age.

Several limitations should be considered. The study population consisted of anonymized residual samples from routine diagnostic testing rather than prospectively recruited healthy individuals. Although samples were selected for complete blood count parameters within the corresponding laboratory reference intervals, detailed clinical information was unavailable, and unrecognized factors affecting erythropoiesis or RBC survival cannot be excluded. The observed distribution should therefore not be interpreted as a formal reference interval. In addition, the protein 4.1a/4.1b ratio was not compared with an independent measurement of RBC survival, and the conversion to erythrocyte life span using the Inaba equation should be considered illustrative rather than a validated individual estimate. Finally, within-person biological variability was not assessed and will require longitudinal measurements in the same individuals.

Overall, direct membrane isolation from small volumes of whole blood substantially simplifies measurement of the protein 4.1a/4.1b ratio and makes this established molecular marker of RBC age suitable for analysis in larger sample series. The present study extends previous experimental applications of protein 4.1a/4.1b by providing, for the first time, an assessment of its interindividual variability in a comparatively large human population. The substantial variation observed despite relatively low analytical variability suggests that the assay captures biological heterogeneity that is not explained by sex or chronological age alone. Establishing the determinants and within-person stability of this variation, and its relationship to independently measured RBC survival, will be important next steps toward evaluating the clinical utility of protein 4.1a/4.1b as a marker of mean RBC age.

## Concluding remarks

We developed a simple and reproducible method for direct isolation of RBC membranes from small volumes of whole blood that enables determination of the protein 4.1a/4.1b ratio without prior RBC washing. The optimized procedure uses 50 μL of whole blood, a volume convenient for routine pipetting and sample processing, while reliable analysis was also possible with substantially smaller volumes when sample availability is limited. Application of the assay to 114 individuals revealed substantial interindividual variation in the protein 4.1a/4.1b ratio, with only modest associations with chronological age and no significant differences between sexes or the investigated age groups. To our knowledge, this represents the first characterization of the population variability of this established molecular marker of RBC age in a comparatively large human cohort. The ability to assess mean RBC age from a single blood sample provides a practical basis for investigating the biological determinants of RBC age and its contribution to laboratory measurements influenced by RBC survival. Further studies should establish within-person biological variability and directly compare the protein 4.1a/4.1b ratio with independent measurements of RBC survival to determine its potential clinical utility.

## Supporting information

Supplementary Code 1: Python script for automated gel densitometry and protein 4.1a/4.1b ratio calculation

