## Supplementary Code 1: Python script for automated gel densitometry and protein 4.1a/4.1b ratio calculation for "A Practical Assay for Assessing Mean Red Blood Cell Age Using the Protein 4.1a/4.1b Ratio": README.txt

SUPPLEMENTARY CODE 1 — GEL-LANE DENSITOMETRIC ANALYSIS
Associated manuscript: A Practical Assay for Assessing Mean Red Blood Cell Age Using the Protein 4.1a/4.1b Ratio

FILES
Supplementary_Code_1.py   Original author-supplied analysis script (renamed only; calculation unchanged)
requirements.txt         Python dependencies
README.txt               This document

PURPOSE AND INPUT
The script processes a CROPPED grayscale-compatible electrophoresis gel image with approximately evenly spaced lanes. It assumes that the target doublet lies ABOVE a separate lower single band in the image. By default it expects six lanes; adjust --lanes if necessary. Dark bands on a light background are treated as positive signal. The lower single band is used to locate lane centers and estimate a common lane width.

INSTALLATION
Use Python 3.10 or newer. In a terminal, navigate to the folder containing these files and run:

    python -m pip install -r requirements.txt

EXAMPLE
    python Supplementary_Code_1.py cropped_gel.png --lanes 6 --outdir results

If the image is vertically inverted, use --flip-vertical. Run:
    python Supplementary_Code_1.py --help
for all command-line options.

OUTPUTS
<image-stem>_results.csv            Numerical measurements and fit diagnostics for each lane
<image-stem>_lane_profiles.png      Baseline-corrected profiles with Gaussian fits
<image-stem>_lane_boundaries.png    Lane boxes and centers overlaid on the input image

QUANTIFICATION DETAILS
The image is converted to grayscale, and dark bands are converted to positive signal. The script averages intensity across each detected lane, removes a baseline using asymmetric least squares (ALS), and smooths the corrected profile. It fits two Gaussians to the upper doublet and one Gaussian to the lower band.

IMPORTANT: The primary reported ratio is NOT the ratio of fitted Gaussian amplitudes or Gaussian areas. Instead, the fitted upper-peak center positions are projected onto the baseline-corrected measured profile, and the profile intensities at those positions are divided:

    upper1_to_upper2_real_curve_peak_ratio = I_corrected(c1) / I_corrected(c2)

The script also exports upper1_to_upper2_gaussian_area_ratio as a separate diagnostic. These two ratios are not interchangeable. The first upper peak is defined by image vertical position, not by automatic protein identification. Users must confirm which peak corresponds to protein 4.1a and which to 4.1b before interpreting the ratio as 4.1a/4.1b.

QUALITY CONTROL AND LIMITATIONS
1. Visually inspect <stem>_lane_boundaries.png to confirm that all lanes are detected and that the selected regions correspond to the intended lanes.
2. Inspect <stem>_lane_profiles.png for correct doublet identification and adequate Gaussian fitting.
3. Review doublet_fit_status; the script still writes a numerical ratio for poorly resolved doublets, so flagged measurements require review rather than automatic acceptance.
4. Check input images for saturation, artifacts, uneven illumination, and unexpected bands. The script does not automatically reject such images.
5. The script assumes lanes are distributed approximately evenly across the cropped image; inappropriate cropping may cause incorrect detection.
6. The --center-half-fraction option is retained for command-line compatibility but is ignored in the current whole-lane implementation.
7. The script does not automatically determine whether upper component 1 or 2 represents protein 4.1a.

REPRODUCIBILITY STATUS
This package contains the supplied analysis code and usage documentation. The code passed Python syntax compilation and its --help command executed. This packaging step did not validate numerical agreement with the study's original gel images or subject-level results; such a comparison is recommended before submission.

DEPENDENCIES
NumPy, pandas, matplotlib, Pillow, and SciPy. Minimum versions are given in requirements.txt; exact versions used for the original analysis were not supplied.
