## Supplementary Code 1: Python script for automated gel densitometry and protein 4.1a/4.1b ratio calculation for "A Practical Assay for Assessing Mean Red Blood Cell Age Using the Protein 4.1a/4.1b Ratio": Supplementary_Code_1.py

```
#!/usr/bin/env python3
"""Quantify six electrophoresis lanes containing an upper doublet and one lower band.

Outputs
-------
<stem>_results.csv            Gaussian parameters, peak areas, and ratios
<stem>_lane_profiles.png      One panel per lane with background-corrected profile and fits
<stem>_lane_boundaries.png    Detected lane positions overlaid on the image

The script treats dark bands as positive signal, detects lane centres and one common lane width from the lower single band, then averages intensity across the complete detected width of each lane,
removes a smooth baseline with asymmetric least-squares (ALS), then fits:
  * upper band region: Gaussian + Gaussian
  * lower band region: one Gaussian

The primary upper-band ratio is obtained by projecting the fitted Gaussian peak\ncentres onto the real background-corrected profile and taking the measured\nprofile heights at those two x positions. Analytic Gaussian areas are retained\nas secondary fit diagnostics.
"""

from __future__ import annotations

import argparse
from pathlib import Path
import warnings

import numpy as np
import pandas as pd
import matplotlib.pyplot as plt
from PIL import Image
from scipy import sparse
from scipy.ndimage import gaussian_filter1d
from scipy.optimize import least_squares
from scipy.signal import find_peaks
from scipy.sparse.linalg import spsolve


def asymmetric_least_squares(y: np.ndarray, lam: float = 2e5,
                             p: float = 0.01, niter: int = 20) -> np.ndarray:
    """Estimate a slowly varying baseline below positive peaks."""
    y = np.asarray(y, dtype=float)
    n = y.size
    d = sparse.diags([1.0, -2.0, 1.0], [0, 1, 2], shape=(n - 2, n))
    penalty = lam * (d.T @ d)
    weights = np.ones(n)
    for _ in range(niter):
        wmat = sparse.spdiags(weights, 0, n, n)
        baseline = spsolve(wmat + penalty, weights * y)
        weights = p * (y > baseline) + (1.0 - p) * (y <= baseline)
    return np.asarray(baseline)


def gaussian(x: np.ndarray, amplitude: float, center: float, sigma: float) -> np.ndarray:
    return amplitude * np.exp(-0.5 * ((x - center) / sigma) ** 2)


def double_gaussian(params: np.ndarray, x: np.ndarray) -> np.ndarray:
    a1, c1, s1, a2, c2, s2 = params
    return gaussian(x, a1, c1, s1) + gaussian(x, a2, c2, s2)


def single_gaussian(params: np.ndarray, x: np.ndarray) -> np.ndarray:
    a, c, s = params
    return gaussian(x, a, c, s)


def fit_doublet(x: np.ndarray, y: np.ndarray, peak_y: int) -> tuple[np.ndarray, float]:
    """Fit two Gaussians using several starting separations; return best fit and RSS."""
    ymax = max(float(np.max(y)), 1e-8)
    xmin, xmax = float(x.min()), float(x.max())
    best = None

    lower = np.array([0.0, xmin, 1.0, 0.0, xmin, 1.0])
    upper = np.array([3*ymax, xmax, 40.0, 3*ymax, xmax, 40.0])

    for sep in (4, 7, 10, 14, 18):
        for frac in (0.35, 0.50, 0.65):
            p0 = np.array([
                ymax * frac, np.clip(peak_y - sep/2, xmin, xmax), 7.0,
                ymax * (1-frac), np.clip(peak_y + sep/2, xmin, xmax), 7.0,
            ])

            def residual(par: np.ndarray) -> np.ndarray:
                r = double_gaussian(par, x) - y
                # Soft penalty discourages numerically identical centers.
                c1, c2 = par[1], par[4]
                separation_penalty = max(0.0, 2.0 - abs(c2-c1)) * ymax
                return np.r_[r, separation_penalty]

            try:
                res = least_squares(residual, p0, bounds=(lower, upper),
                                    max_nfev=10000)
            except ValueError:
                continue
            rss = float(np.sum((double_gaussian(res.x, x) - y) ** 2))
            if best is None or rss < best[1]:
                best = (res.x, rss)

    if best is None:
        raise RuntimeError("Double-Gaussian fitting failed")

    par, rss = best
    # Sort by migration coordinate: upper component first, lower component second.
    first = par[:3]
    second = par[3:]
    if first[1] > second[1]:
        par = np.r_[second, first]
    return par, rss


def fit_single(x: np.ndarray, y: np.ndarray, peak_y: int) -> tuple[np.ndarray, float]:
    ymax = max(float(np.max(y)), 1e-8)
    p0 = np.array([ymax, float(peak_y), 8.0])
    lower = np.array([0.0, float(x.min()), 1.0])
    upper = np.array([3*ymax, float(x.max()), 45.0])
    res = least_squares(lambda p: single_gaussian(p, x)-y, p0,
                        bounds=(lower, upper), max_nfev=10000)
    rss = float(np.sum((single_gaussian(res.x, x)-y)**2))
    return res.x, rss


def detect_lane_intervals(signal: np.ndarray, n_lanes: int = 6,
                          center_half_fraction: float = 0.20
                          ) -> tuple[list[tuple[int, int]], list[tuple[int, int]], np.ndarray]:
    """Detect lanes from the sharper lower single band and apply one common width.

    Workflow
    --------
    1. Find the lower single-band region from the vertical intensity profile.
    2. Build a horizontal intensity profile using only a narrow strip around
       that lower band.
    3. Detect exactly one lane centre inside each horizontal sector.
    4. Measure each lane's horizontal width at half prominence.
    5. Use the median measured width as one common width for all lanes.
    6. Re-centre each common-width box on the local half-prominence midpoint.

    The actual electrophoresis profile is still calculated from the complete
    height of each lane rectangle. The lower band is used only to define lane
    centres and lane width.
    """
    from scipy.signal import peak_widths

    h, w = signal.shape

    # 1. Find the lower single band vertically
    vertical_profile = np.mean(signal, axis=1)
    vertical_smooth = gaussian_filter1d(vertical_profile, sigma=3.0)
    search_start = int(round(0.45 * h))
    lower_y = search_start + int(np.argmax(vertical_smooth[search_start:]))

    # 2. Horizontal profile from a narrow strip around the lower band
    strip_half_height = max(5, int(round(0.08 * h)))
    y0 = max(0, lower_y - strip_half_height)
    y1 = min(h, lower_y + strip_half_height + 1)

    x_profile = np.mean(signal[y0:y1, :], axis=0)
    smooth = gaussian_filter1d(x_profile, sigma=max(4.0, w / 250.0))
    slow_background = gaussian_filter1d(x_profile, sigma=max(35.0, w / 18.0))
    corrected = smooth - slow_background

    # 3. Detect exactly one lane peak in each horizontal sector
    sector_edges = np.linspace(0, w, n_lanes + 1).astype(int)
    peak_positions = []
    for i in range(n_lanes):
        left = int(sector_edges[i])
        right = int(sector_edges[i + 1])
        margin = max(2, int(round(0.10 * (right - left))))
        lo = left + margin
        hi = right - margin
        if hi <= lo:
            lo, hi = left, right
        local = corrected[lo:hi]
        if local.size == 0:
            peak = int(round(0.5 * (left + right)))
        else:
            peak = lo + int(np.argmax(local))
        peak_positions.append(peak)

    peak_positions = np.asarray(peak_positions, dtype=int)
    spacing = float(np.median(np.diff(peak_positions))) if n_lanes > 1 else float(w)

    # 4. Measure actual horizontal widths of the lower bands
    widths, _, _, _ = peak_widths(corrected, peak_positions, rel_height=0.50)
    valid = (
        np.isfinite(widths)
        & (widths > 0.20 * spacing)
        & (widths < 0.85 * spacing)
    )

    if np.any(valid):
        measured_common_width = float(np.median(widths[valid]))
    else:
        measured_common_width = 0.55 * spacing

    # Expand beyond FWHM so the full visible lane is included, while preventing overlap.
    common_width = 1.20 * measured_common_width
    common_width = max(0.35 * spacing, common_width)
    common_width = min(common_width, 0.75 * spacing)
    half_width = max(3, int(round(0.5 * common_width)))

    # 5. Refine each centre from the local lower-band edges
    centers = []
    search_radius = max(6, int(round(0.35 * spacing)))

    for peak in peak_positions:
        lo = max(1, int(peak - search_radius))
        hi = min(w - 1, int(peak + search_radius + 1))
        if hi <= lo:
            centers.append(float(peak))
            continue

        local_peak = lo + int(np.argmax(corrected[lo:hi]))
        ww, _, ll, rr = peak_widths(corrected, [local_peak], rel_height=0.50)

        width_ok = (
            np.isfinite(ww[0])
            and np.isfinite(ll[0])
            and np.isfinite(rr[0])
            and ww[0] > 0.20 * spacing
            and ww[0] < 0.85 * spacing
        )

        if width_ok:
            center = 0.5 * (float(ll[0]) + float(rr[0]))
        else:
            local = corrected[lo:hi]
            baseline = float(np.percentile(local, 20))
            weights = np.clip(local - baseline, 0.0, None)
            if np.sum(weights) > 0:
                x = np.arange(lo, hi, dtype=float)
                center = float(np.sum(x * weights) / np.sum(weights))
            else:
                center = float(local_peak)
        centers.append(center)

    centers = np.asarray(centers, dtype=float)

    # 6. Apply exactly the same measured width to every lane
    intervals = []
    for center in centers:
        c = int(round(center))
        x0 = max(0, c - half_width)
        x1 = min(w, c + half_width)
        intervals.append((x0, x1))

    full_intervals = list(intervals)
    return intervals, full_intervals, centers

def choose_band_windows(profile: np.ndarray) -> tuple[tuple[int, int], tuple[int, int], int, int]:
    """Identify the two main vertical band groups and define non-overlapping windows."""
    n = len(profile)
    sm = gaussian_filter1d(profile, 3)
    peaks, _ = find_peaks(sm, distance=max(12, n//8), prominence=max(np.ptp(sm)*0.05, 1e-5))
    if len(peaks) >= 2:
        strongest = peaks[np.argsort(sm[peaks])[-2:]]
        p_top, p_bottom = sorted(strongest)
    else:
        p_top, p_bottom = int(0.27*n), int(0.70*n)

    midpoint = (p_top+p_bottom)//2
    margin = max(4, int(0.04*n))
    top_window = (max(0, p_top-int(0.24*n)), min(midpoint-margin, p_top+int(0.24*n)))
    bottom_window = (max(midpoint+margin, p_bottom-int(0.22*n)), min(n, p_bottom+int(0.22*n)))
    return top_window, bottom_window, p_top, p_bottom


def analyze(image_path: Path, outdir: Path, n_lanes: int, center_half_fraction: float,
            baseline_lambda: float, baseline_p: float, flip_vertical: bool) -> None:
    image = np.asarray(Image.open(image_path).convert("L"), dtype=float)/255.0
    if flip_vertical:
        image = image[::-1, :]
    signal = 1.0-image  # dark bands become positive
    h, w = signal.shape

    intervals, full_intervals, centers = detect_lane_intervals(signal, n_lanes, center_half_fraction)
    outdir.mkdir(parents=True, exist_ok=True)
    stem = image_path.stem

    # Boundary check image
    fig, ax = plt.subplots(figsize=(14, 3.2))
    ax.imshow(image, cmap="gray", aspect="auto", origin="upper")
    from matplotlib.patches import Rectangle
    for i, ((x0, x1), (f0, f1), c) in enumerate(zip(intervals, full_intervals, centers), start=1):
        # Solid rectangle: complete common-width lane area used for analysis.
        ax.add_patch(Rectangle((x0, 0), x1-x0, h-1, fill=False,
                               linestyle="-", linewidth=2.0))
        ax.axvline(c, linestyle="--", linewidth=1.0, alpha=0.8)
        ax.text(c, 8, str(i), ha="center", va="top",
                bbox=dict(facecolor="white", alpha=0.70, edgecolor="none"))
    ax.set_title("Whole-lane analysis: lane centres/width from lower single band")
    ax.set_xlabel("x (pixels)")
    ax.set_ylabel("migration coordinate y (pixels)")
    fig.tight_layout()
    fig.savefig(outdir/f"{stem}_lane_boundaries.png", dpi=200)
    plt.close(fig)

    ncols = 2
    nrows = int(np.ceil(n_lanes/ncols))
    fig, axes = plt.subplots(nrows, ncols, figsize=(13, 3.5*nrows), squeeze=False)
    rows = []

    for lane_idx, (x0, x1) in enumerate(intervals, start=1):
        raw = np.mean(signal[:, x0:x1], axis=1)
        baseline = asymmetric_least_squares(raw, lam=baseline_lambda, p=baseline_p)
        corrected = np.clip(raw-baseline, 0, None)
        corrected = gaussian_filter1d(corrected, sigma=1.0)

        top_win, bottom_win, p_top, p_bottom = choose_band_windows(corrected)
        tx = np.arange(top_win[0], top_win[1], dtype=float)
        bx = np.arange(bottom_win[0], bottom_win[1], dtype=float)
        ty = corrected[top_win[0]:top_win[1]]
        by = corrected[bottom_win[0]:bottom_win[1]]

        top_par, top_rss = fit_doublet(tx, ty, p_top)
        lower_par, lower_rss = fit_single(bx, by, p_bottom)
        a1, c1, s1, a2, c2, s2 = top_par
        a3, c3, s3 = lower_par
        area1 = a1*s1*np.sqrt(2*np.pi)
        area2 = a2*s2*np.sqrt(2*np.pi)
        area3 = a3*s3*np.sqrt(2*np.pi)
        eps = np.finfo(float).eps

        # Project the fitted Gaussian maxima (their center coordinates) onto
        # the REAL measured, background-corrected profile. The resulting y
        # values are the experimental peak heights used for the main ratio.
        real_height1 = float(np.interp(c1, tx, ty))
        real_height2 = float(np.interp(c2, tx, ty))
        real_peak_ratio = real_height1 / (real_height2 + eps)

        separation = abs(c2-c1)
        resolution = separation/(2*(s1+s2))  # approximate chromatographic resolution
        status = "OK" if resolution >= 0.5 else "poorly resolved; interpret doublet cautiously"

        rows.append({
            "lane": lane_idx, "x_start_px": x0, "x_end_px": x1,
            "upper_1_center_px": c1, "upper_1_amplitude": a1,
            "upper_1_sigma_px": s1, "upper_1_area": area1,
            "upper_2_center_px": c2, "upper_2_amplitude": a2,
            "upper_2_sigma_px": s2, "upper_2_area": area2,
            "lower_center_px": c3, "lower_amplitude": a3,
            "lower_sigma_px": s3, "lower_area": area3,
            "upper_1_real_curve_height_at_gaussian_center": real_height1,
            "upper_2_real_curve_height_at_gaussian_center": real_height2,
            "upper1_to_upper2_real_curve_peak_ratio": real_peak_ratio,
            "upper1_to_upper2_gaussian_area_ratio": area1/(area2+eps),
            "upper1_fraction_of_doublet": area1/(area1+area2+eps),
            "upper2_fraction_of_doublet": area2/(area1+area2+eps),
            "doublet_to_lower_area_ratio": (area1+area2)/(area3+eps),
            "upper1_to_lower_area_ratio": area1/(area3+eps),
            "upper2_to_lower_area_ratio": area2/(area3+eps),
            "doublet_center_separation_px": separation,
            "doublet_resolution_approx": resolution,
            "doublet_fit_status": status,
            "doublet_fit_rss": top_rss, "lower_fit_rss": lower_rss,
        })

        ax = axes[(lane_idx-1)//ncols, (lane_idx-1)%ncols]
        yy = np.arange(h)
        ax.plot(yy, corrected, label="background-corrected profile", linewidth=1.4)
        ax.plot(tx, double_gaussian(top_par, tx), label="double-Gaussian sum", linewidth=2)
        ax.plot(tx, gaussian(tx, a1, c1, s1), linestyle="--", label="upper Gaussian 1")
        ax.plot(tx, gaussian(tx, a2, c2, s2), linestyle="--", label="upper Gaussian 2")
        ax.scatter([c1, c2], [real_height1, real_height2], s=35, zorder=5,
                   label="Gaussian centers projected to real curve")
        ax.vlines([c1, c2], 0, [real_height1, real_height2],
                  linestyles=":", linewidth=1.1)
        ax.plot(bx, single_gaussian(lower_par, bx), label="lower Gaussian", linewidth=2)
        ax.axvline(c1, linewidth=0.8)
        ax.axvline(c2, linewidth=0.8)
        ax.axvline(c3, linewidth=0.8)
        ax.set_title(f"Lane {lane_idx}: real-curve peak ratio U1/U2={real_peak_ratio:.3f}")
        ax.set_xlabel("migration coordinate y (pixels)")
        ax.set_ylabel("corrected optical-density signal (a.u.)")
        ax.grid(alpha=0.2)
        if lane_idx == 1:
            ax.legend(fontsize=8, ncol=2)

    for j in range(n_lanes, nrows*ncols):
        axes[j//ncols, j%ncols].axis("off")
    fig.tight_layout()
    fig.savefig(outdir/f"{stem}_lane_profiles.png", dpi=220)
    plt.close(fig)

    results = pd.DataFrame(rows)
    results.to_csv(outdir/f"{stem}_results.csv", index=False, float_format="%.8g")
    pd.set_option("display.max_columns", None)
    print(results[["lane",
                   "upper_1_real_curve_height_at_gaussian_center",
                   "upper_2_real_curve_height_at_gaussian_center",
                   "upper1_to_upper2_real_curve_peak_ratio",
                   "upper1_to_upper2_gaussian_area_ratio",
                   "doublet_to_lower_area_ratio",
                   "doublet_resolution_approx", "doublet_fit_status"]].to_string(index=False))
    print(f"\nSaved results in: {outdir.resolve()}")


def main() -> None:
    parser = argparse.ArgumentParser(description=__doc__,
                                     formatter_class=argparse.RawDescriptionHelpFormatter)
    parser.add_argument("image", type=Path, help="cropped gel image containing the lanes")
    parser.add_argument("--outdir", type=Path, default=Path("gel_analysis"))
    parser.add_argument("--lanes", type=int, default=6)
    parser.add_argument("--center-half-fraction", type=float, default=0.20,
                        help="retained only for command compatibility; ignored in whole-lane mode; "
                             "0.20 means the central 40%% is analyzed (default: 0.20)")
    parser.add_argument("--baseline-lambda", type=float, default=2e5,
                        help="ALS smoothness; larger gives smoother baseline")
    parser.add_argument("--baseline-p", type=float, default=0.01,
                        help="ALS asymmetry for positive peaks (default: 0.01)")
    parser.add_argument("--flip-vertical", action="store_true",
                        help="reverse vertical migration direction before analysis")
    args = parser.parse_args()
    analyze(args.image, args.outdir, args.lanes, args.center_half_fraction,
            args.baseline_lambda, args.baseline_p, args.flip_vertical)


if __name__ == "__main__":
    main()
```
